# Field and genetic evidence support the photosynthetic performance index (PI_ABS_) as an indicator of rice grain yield

**DOI:** 10.1101/2023.02.08.527648

**Authors:** Andrés Alberto Rodríguez, Juan Manuel Vilas, Gustavo Daniel Sartore, Rodolfo Bezus, José Colazo, Santiago Javier Maiale

## Abstract

The effective increase of the rice breeding process for grain yield could be sustained by developing efficient tools to accelerate plant selection through the rapid determination of reliable predictors. Here, we have described different associations between grain yield and photosynthetic parameters simply and fast obtainable by a non-invasive technique in flag leaf during the anthesis stage. Among the analyzed photosynthetic parameters, the photosynthetic performance index (PI_ABS_) stood out for its strong association with grain yield. A genome-wide association analysis determined in plants from a rice diversity panel at tillering stage indicated the presence of a quantitative trait locus on chromosome 9 characterized by a set of candidate chloroplastic genes with contrasting haplotypes for PI_ABS_. An analysis of these haplotypes indicated a separation into two groups. One with haplotypes linked to high values of PI_ABS_, which were associated almost exclusively with *Japonica* spp. subpopulations, and another with haplotypes linked to low values of PI_ABS_, which were associated exclusively with *Indica* spp. subpopulations. Genotypes of the *Japonica* spp. subpopulations showed high values in panicle weight, a yield components parameter, compared with the *Indica* spp. subpopulations genotypes. The results of this work suggested that PI_ABS_ could be an early predictor of grain yield at the tillering stage in rice breeding processes.

## 1. Introduction

Current evidence shows that there is a genetic background in the main cultivated species to improve the sustainability of their crops by increasing their yields through an improvement in the environmental adaptation of plants to the environments in which they are grown. However, the screening for higher yields under field conditions often imposes troubles due to the variability of climate, edaphic conditions and management practices (Tavakkoli et al., 2012). In the case of the rice species *Oryza sativa*, there is also the fact that its genotypes differ enormously in the levels of grain yield owing to the vast diversity of genetic constitutions (Xing and Zhang, 2010). The use of indirect selection criteria in breeding for better yields was reported in rice (Shen et al., 2001). However, for indirect selection to be successful, the specific traits must have a high correlation with yield (Richards et al., 2001). Therefore, developing robust physiological indicators of easy and rapid determination that correlate with yield parameters in the field could be a practical indirect approach for mass screening populations as significant as used in breeding programs.

Field experiments have shown that high yield in rice is causally related to the protection of the photosynthetic apparatus through excess light dissipation (Wang et al., 2014). The OJIP test is a technique that depicts different OJIP parameters linked with the state of structures and functionalities related to the photosystem II (PSII) complex (Calzadilla et al., 2022). It includes functional parameters such as F_V_/F_M_ and ΨE_0_ and structural parameter as γRC. F_V_/F_M_ represents the maximum photochemical efficiency of PSII, which represents the maximum efficiency with which an absorbed photon result in the reduction of the quinone A. ΨE_0_ represents the maximum efficiency of electron transport in PSII beyond reduced quinone A, and γRC reflects alterations in the density of active PSII reaction centre (RC) through changes in the active chlorophyll associated to the RC (Strasser et al., 2000). Besides, Živčák et al. (2008) reported the photosynthetic performance index (PI_ABS_) derived from the union of γRC, F_V_/F_M_ and ΨE_0_ in a single equation PI_ABS_ = (γRC / 1 - γRC). (F_V_/F_M_ / 1 - F_V_/F_M_). (ΨE_0_ / 1 - ΨE_0_). These authors described the PI_ABS_ as an integrative and sensitive parameter to register the combined changes of these last three OJIP parameters in a unique value of plant vitality. Some authors have shown that super-high-yielding hybrid rice flag leaves are causally related to PI_ABS_ (Zhang et al., 2015). Parent’s election in breeding programs depends on screening and selecting genotypes with better performance characteristics. However, for selective breeding to be practical, genetic variation must be present in the screened population (Hill, 2001). In the case of *O. sativa*, the genetic diversity is encompassed by two subspecies and five subpopulations. The subspecies *Japonica ssp*. groups include the subpopulations temperate Japonica, aromatic and tropical Japonica and the subspecies *Indica ssp*. groups include the subpopulations indica and aus (Garris et al., 2005). These five subpopulations are well represented in the rice diversity germplasm collection called Rice Diversity Panel 1 (RDP1), which contains 413 genotypes. The RDP1 spans a wide genetic variability based on the diversity of environmental growth conditions at the origin and breeding histories of *O. sativa* accessions (Zhao et al., 2011).

In order to evaluate potential physiological parameters as indirect yield indicators with easy and rapid determination related to structural and functional parameters linked to PSII, we performed two experiments (Experiments 1 and 2). Experiment 1 was performed to characterize the relationship between yield parameters with structures and functionalities of the PSII determined at anthesis in the flag leaf of rice genotypes with contrasting grain yield in the field plot. In Experiment 2, different genome-wide association analysis (GWAS) was performed using a phenotypification determined at the tillering stage in a leaf of plants from RDP1 genotypes based on photosynthetic parameters, particularly PI_ABS_. This last experiment was designed to generate meaningful information about the connections of the rice population structure with the relationship between yield and the PSII performance and with the PI_ABS_ capacity as an early predictor of grain yield during the tillering stage in rice breeding processes.

## 2. Materials and Methods

### 2.1. Plant material and growth conditions

Seeds of 10 rice genotypes with contrasting grain yields were used in Experiment 1. The selected genotypes were H458, H426-1-1-1, Don Ignacio, H294, R-03, Don Justo, H426-25, Yerua and H420. The selected genotypes were the cultivars Don Ignacio FCAyF, Don Justo FCAyF and Yerúa and the inbreed lines H458, H426-1-1, H294, H426-25, R-03 and H420, all belonging to FCAyF rice breeding program. Seeds from all genotypes were mechanically sown in plots of 20 square meters in a randomized design with three replications with 20 cm in the space between rows and with a density of 300 seeds m^-2^. The amounts of fertilizer applied as a basal dressing were 6.84 g N m^-2^, 2.7 g P m^-2^ and 8.5 g Km^-2^. The trial was conducted under flooding until the maturity stage.

Seeds of 283 genotypes were used in Experiment 2, these comprising 45 aus, 59 indica, 58 temperate and 76 tropical japonica, eight aromatic and 37 admixed rice genotypes. Seeds from all genotypes were sown in Petri dishes on two layers of Whatman Nº 5 filter paper, rinsed with 7 mL carbendazim 0.025 %p/v and incubated at 30 °C in darkness for three days until germination. Each resultant seedling was transplanted into a plastic pot (10 L) containing sterilized organic soil extract as substrate. The pots were transferred to field environmental conditions into a plot in a completely randomized design with three replications, and the trial was conducted under flooding until the maturity stage.

### 2.2. Determination of photosynthetic parameters

The photosynthetic parameters were determined by analyzing the chlorophyll fluorescence emission kinetics according to Gazquez et al. (2015). A portable fluorometer was used, HANDY PEA (Hansatech Instruments® Ltd., King’s Lynn, Norfolk, UK). Briefly, blade sections of intact leaves were covered with a leaf clip to adapt them to darkness for 20 min. The flag leaf represented the intact leaves at the anthesis stage (Experiment 1) or the uppermost fully expanded leaf in 11-week-old plants at tillering stage (Experiment 2). Then, the blade sections were exposed to a 3 s pulse of red light (650 nm, 3500 μmol photons m^-2^ s^-1^). The raw fluorescence data of the fluorescence emission kinetic was processed by the PEA plus software (Hansatech Instrument, UK) to determine the different OJIP parameters. The OJIP parameters described above, γRC, F_V_/F_M_ and ΨE_0_, were calculated according to the equations described by Puig et al. (2021).

### 2.3. Determination of yield parameters

In all experiments, plants were harvested, panicles were threshed manually, and the grain was dried in an oven at 40 ºC until 14% humidity. For Experiment 1, the yield parameters were determined per plot as the number of grains per square meter (Grains. m^-2^) and the Kg of grains per ha^-1^ (Y). For Experiment 2, the number of panicles (NP) and the weight of filled grains (WFG) were determined per plant at harvest. Then, the panicle weight (PW) was calculated as PW = WFG. NP^-1^.

### 2.4. Genome-wide association and linkage disequilibrium analyses

The GWAS studies and the linkage disequilibrium (LD) were performed based on the HDRA dataset (HDRA6.4) genotyping from the RDP1 genotypes consisting of 700,000 single nucleotide polymorphism (SNPs) described by McCouch et al. (2016). Three replicates of the complete set of samples were used to perform the phenotyping data for each trait. GWAS mapping was performed considering settings and recommendations described by McCouch et al. (2016) using the Tassel software (Tassel v5.0). Briefly, GWAS was running using a linear mixed model in the EMMAX algorithm (Kang et al., 2010), which considers the underlying population structure by including a kinship matrix as a covariate. A minor allele frequency (MAF) threshold of 0.05 (MAF < 0.05) was applied to discard markers with exceedingly rare alleles. When the GWAS were run across all available RDP1 genotypes, SNPs at MAF > 0.05 in individual subpopulations were combined. Then, three additional principal component covariates, derived from a principal component analysis (PCA) to characterize the RDP1 genetic structure, were added to the model. Based on the approximate significance value where the observed p-values exceed the expected number in the Q-Q plots, a significance threshold of 10^−4^ was used to identify significant SNP across all analyses. The quantitative trait locus (QTL) determined in the GWAS peak on chromosome 9 were delimited according to McCouch et al. (2016). Regions with significant SNPs were identified as having three or more significant SNPs in a 200 kb region. They overlapped when they shared significant SNPs to form a single QTL region. GWAS LD among markers on chromosome 9 was calculated using pairwise r square between SNPs and the LD analysis function with an LD windows size of 500 SNPs in the Tassel software.

### 2.5. Gene targeting, gene annotation and haplotype analyses

The genes within the QTL regions were identified using the data of the RICEBASE (www.ricebase.org) database and a protein subcellular localization prediction analysis. The protein sequence of each gene was downloaded from the MSU Rice Genome Annotation Project version 7. The protein subcellular localization prediction analysis was performed, integrating all information of these protein sequences from the databases: WoLF PSORT, Plant-mPLoc and TargetP. Gene annotation about the GWAS peak genes was performed, integrating all information from the databases: Gramene, Uniprot and KEEG. Haplotypes from all genes with the predicted chloroplastic location were collected and entered along with their associated PI_ABS_ average values into statistical analysis software to select genes with contrasting haplotypes in PI_ABS_.

### 2.6. Statistics

All data sets were tested for normality. Data from photosynthetic and yield parameters, including the PI_ABS_ average values associated with haplotypes of genes with contrasting haplotypes in PI_ABS_, were subjected to ANOVA and post hoc analyses DGC tests (Di Rienzo, Guzmán, Casanoves, 2002) and to Student’s t-test. These data were also subjected to linear correlation analyses, principal component analyses (PCA), and some linear regression analyses using the Infostat® statistical software package used throughout the study (Di Rienzo et al., 2018).

## 3. Results

### 3.1. Experiment 1. Relationship between the yield with structures and functionalities of the photosystem II in the flag leaf of contrasting rice genotypes in grain yield

A PCA showed that the variability associated with the set of all parameters separated the data of genotypes with above-average yield or high yield (HY, 10 Tn/ha on average) from genotypes with below-average yield or typical yield (NY, 8.3 Tn/ha in average). The data separation was due to PC1, which retained 71.3% of the total dataset variability (Fig. 1). The reported eigenvectors associated with each original variable weighted to form PC1 indicated PI_ABS_, ΨE_O_, F_V_/F_M_, γRC and Kg of grains. m^-2^ (Y) received the highest weights (Table S1). On the other hand, Y explained 18.5% of the variability retained by PC2. Therefore, these results suggested that multiple parameters related to the PSII and mainly Y explained the distinction between HY and NY genotypes mainly by the variability explained by PC1.

**FIGURE 1.**
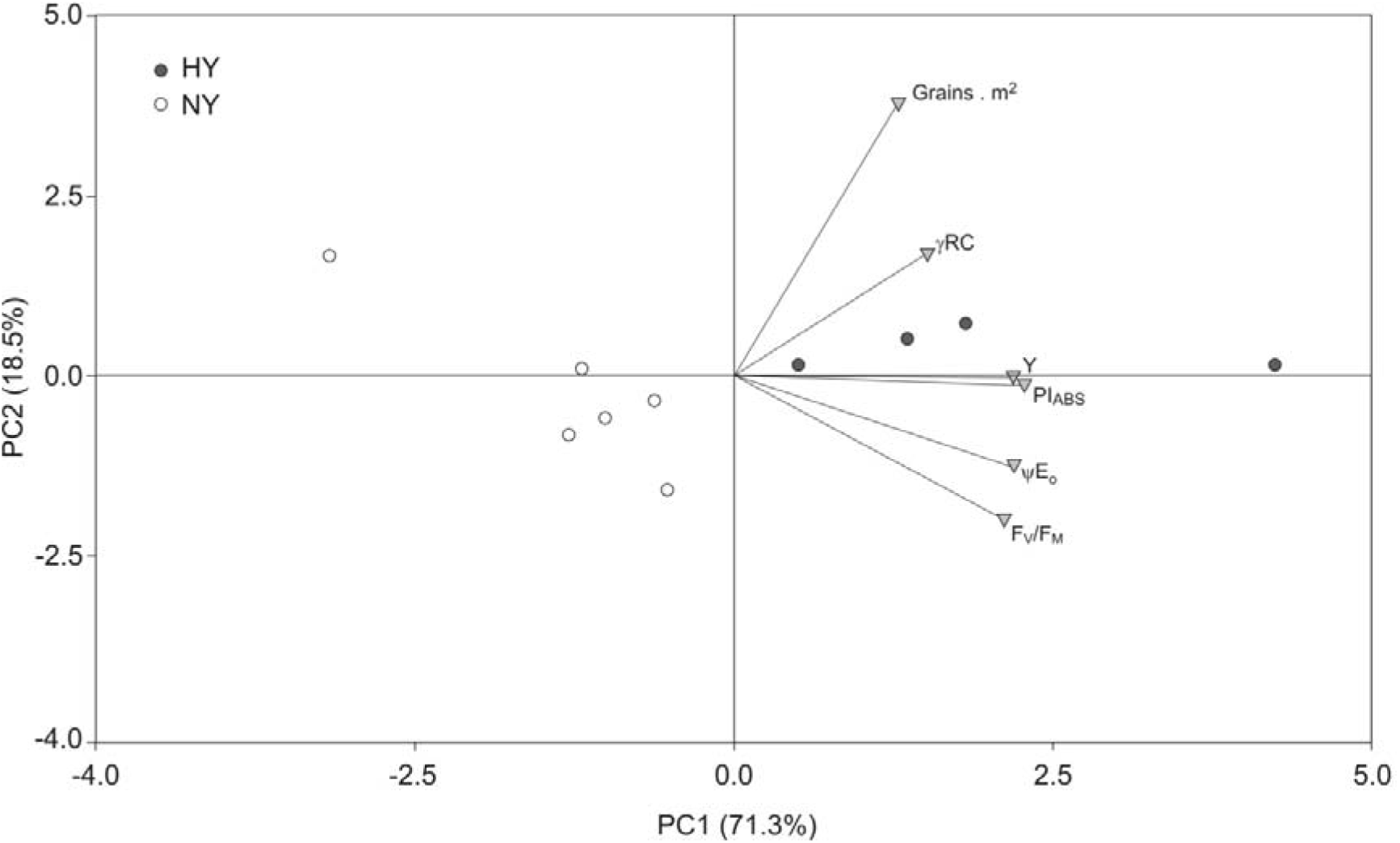
PCA of yield and OJIP parameters data from genotypes with contrasting in grain yield at anthesis stage. HY, genotypes with high yield; NY, genotypes with normal yield. Yield parameters: Grains. m^-2^, number of grains per m^-2^; Y, Kg of grains. ha^-1^. OJIP parameters: F_V_/F_M_, maximum photochemical efficiency of PSII; ΨE_0_, maximum efficiency of electron transport in PSII beyond reduced quinone A; γRC, density of active PSII reaction centre; PI_ABS_, performance index.

In addition, the PCA results revealed multiple associations between yield and PSII parameters. Linear correlation analysis indicated many significantly positive correlations among structure and functional parameters of the PSII in flag leaf with yield parameters (Table 1). The most relevant results were the high correlations between the Y with all PSII parameters, particularly with PI_ABS_ and ΨE_O_ (r = 0.85 in both cases). This data was in line with another statistical analysis in which a complementary model from different linear regression analyses indicated that 72% of the variation of Y was explained by PI_ABS_ (Fig. 2). All these statistical results suggested a relationship between yield and PSII parameters in which PI_ABS_ explained most of the contrast observed.

**Table 1.**
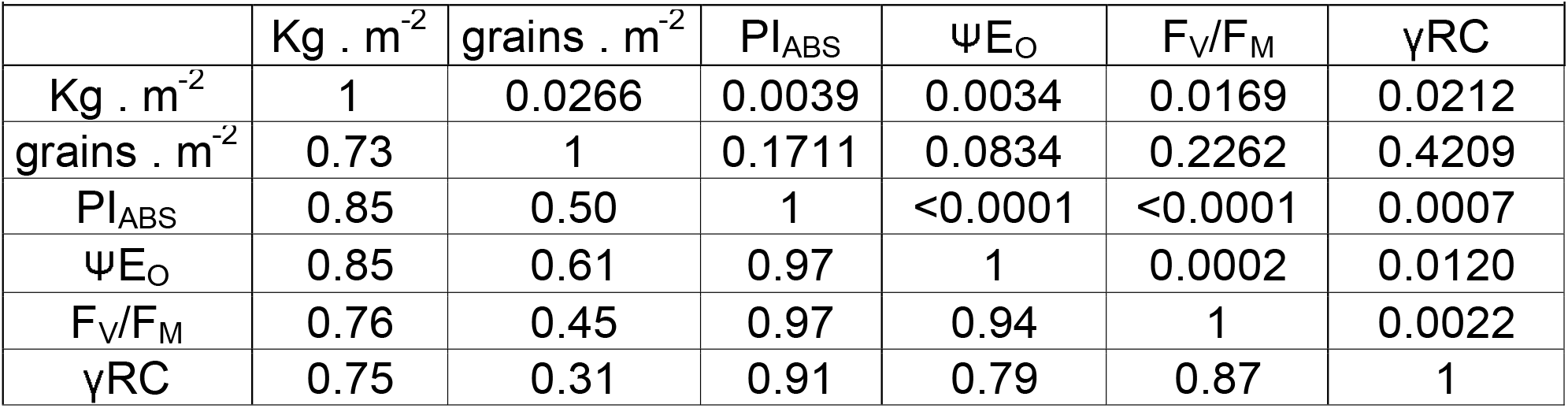
Correlation matrix of structure and functional parameters of the photosynthetic apparatus in rice flag leaf with yield parameters. The matrix correlation presents the Pearson r values and its corresponding P values to each correlation between the different parameters.

**FIGURE 2.**
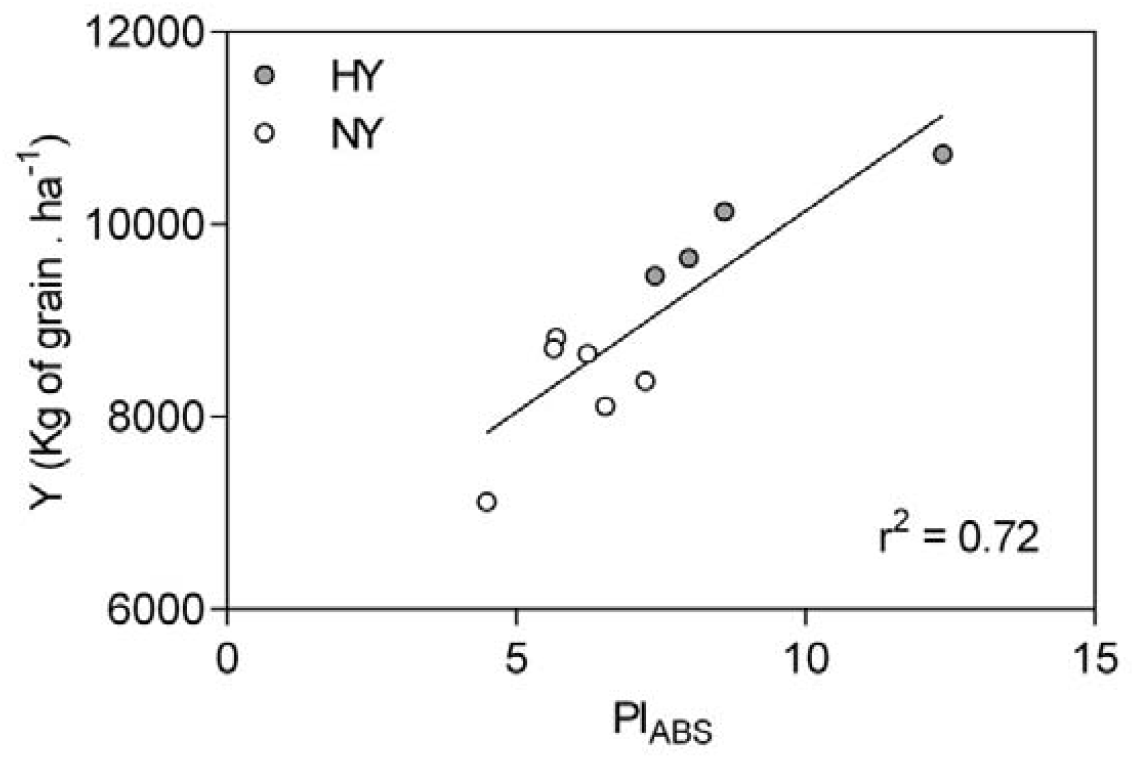
Linear regression analysis of PI_ABS_ and Y data from genotypes with contrasting in grain yield at the anthesis stage. The model’s goodness of fit was determined by the coefficient of determination R square (r^2^). The solid line in each graph represents the best regression linear fit model between dependent (Y-axis) and independent (X-axis) variables.

### 3.2. Experiment 2. Genome-wide association analysis using an OJIP phenotypification in plants of RDP1 genotypes

A GWAS performed using a phenotypification by PI_ABS_ determination in plants of genotypes from the RDP1 rice diversity panel indicated the presence of multiple significant molecular markers grouped in a GWAS peak on chromosome 9 that involved 170 genes (Fig. 3A). Other complementary GWAS performed using phenotypifications by ΨE_O_, and γRC determinations also indicated multiple significant molecular markers in common with PI_ABS_ in this GWAS peak on chromosome 9 (Fig. 3A). A posterior analysis showed that 59 of these genes had a predicted location in the chloroplast (Fig. 3B) and were in regions with high LD values (Fig. 3C). Twenty per cent of genes found in this work could not be characterized, and the remaining 80% were proteins related to functions in the chloroplastid (Table 2). In addition, there were twenty chloroplastic genes with contrasting haplotypes in PI_ABS_ (CGPI_ABS_, Fig. 3D).

**FIGURE 3.**
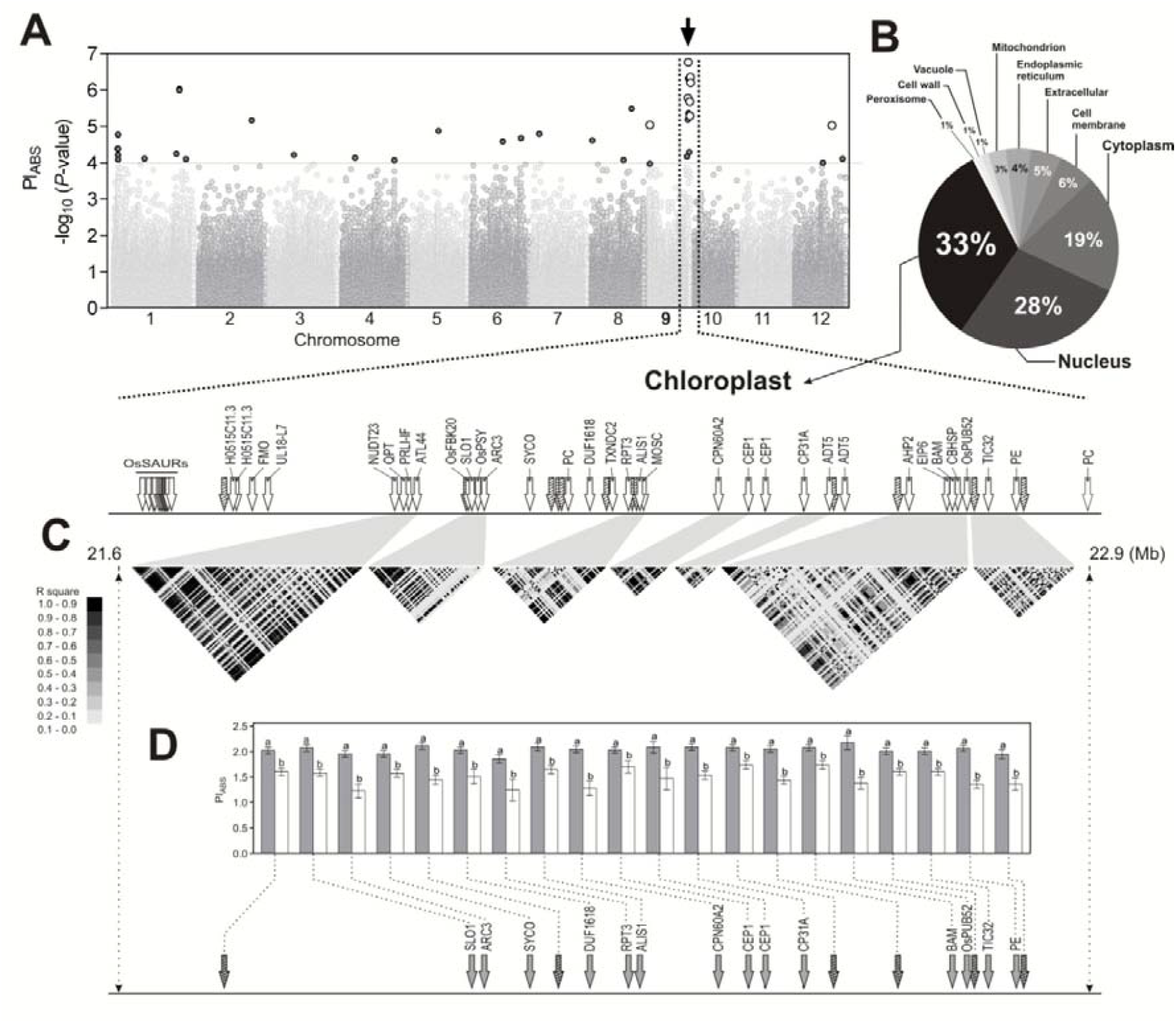
GWAS using PI_ABS_ determined in plants of the rice diversity panel RDP1 at tillering stage. **A**. Manhattan plot using the RDP1 genotyping and the PI_ABS_ phenotyping. The x-axis shows the SNPs along each chromosome; the y-axis is the - log_10_ (*p*-value) for the association. Gray and white circles above dotted line represent significant SNPs with -log_10_(*p*-value) > 4. The gray circles correspond to SNPs registered in GWAS with PI_ABS_ phenotyping. The white circles correspond to significant SNPs registered in GWAS with PI_ABS_ phenotyping but also with ΨE_0_ and γRC phenotyping. **B**. Pie chart with results of gene percentages for each subcellular location obtained by a protein subcellular localization prediction analysis. **C**. QTL region in the GWAS peak on chromosome 9 enrichment with chloroplastic genes. The horizontal bar shows a physical map of chromosome 9 (21.6–22.9 Mb); the chloroplastic genes with associated or uncharacterized functions were represented by unfilled or filled arrows, respectively. **D**. The main haplotypes associated to from candidate chloroplastic genes with contrasting haplotypes for PI_ABS_ were analyzed. Grey and white bars represent haplotypes associated with PI_ABS_ average high and low values, respectively (ANOVA and post hoc analysis DGC tests *p* < 0.05, n = 5 per genotype; Different letters represent significant differences between haplotypes).

**Table 2.**
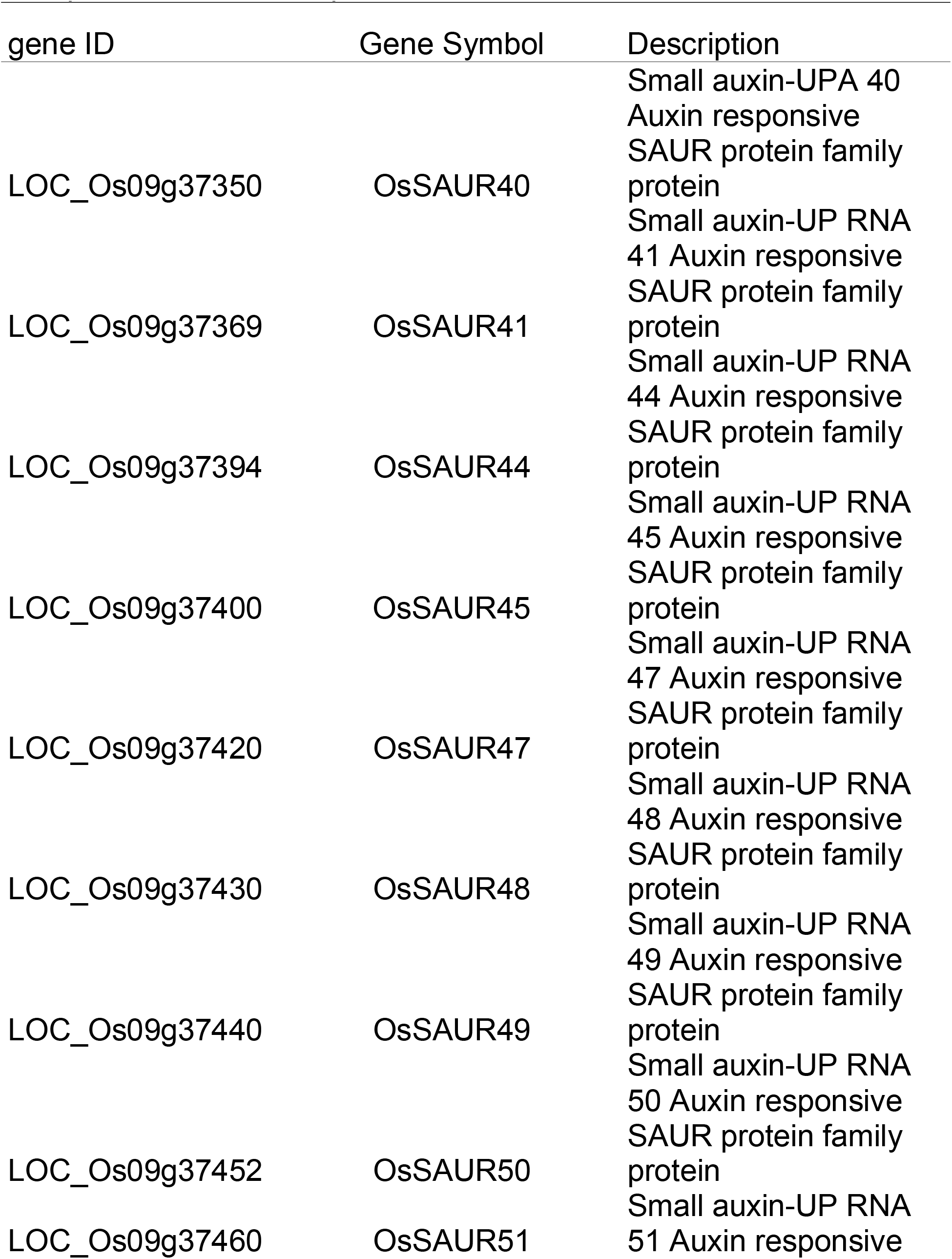
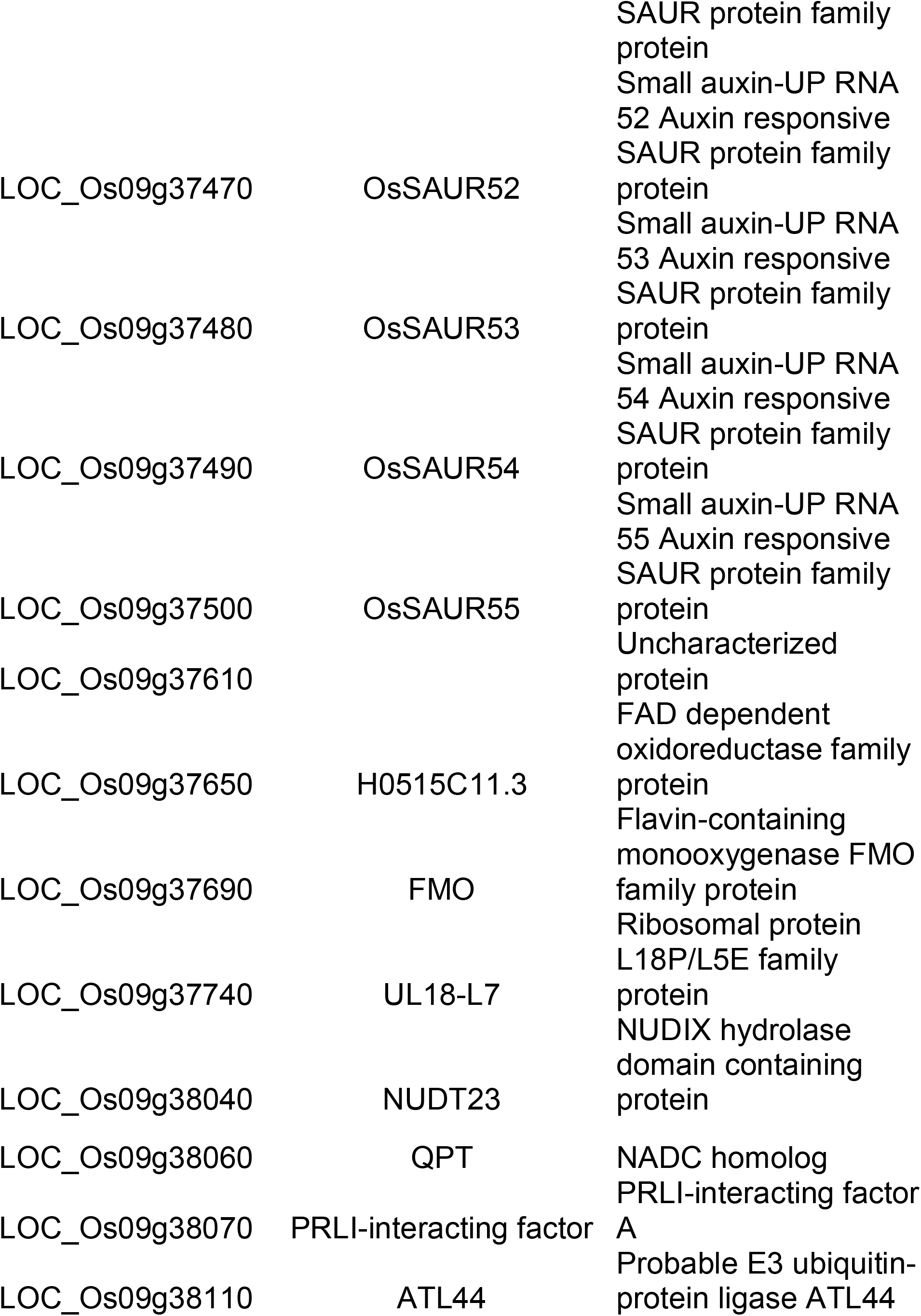
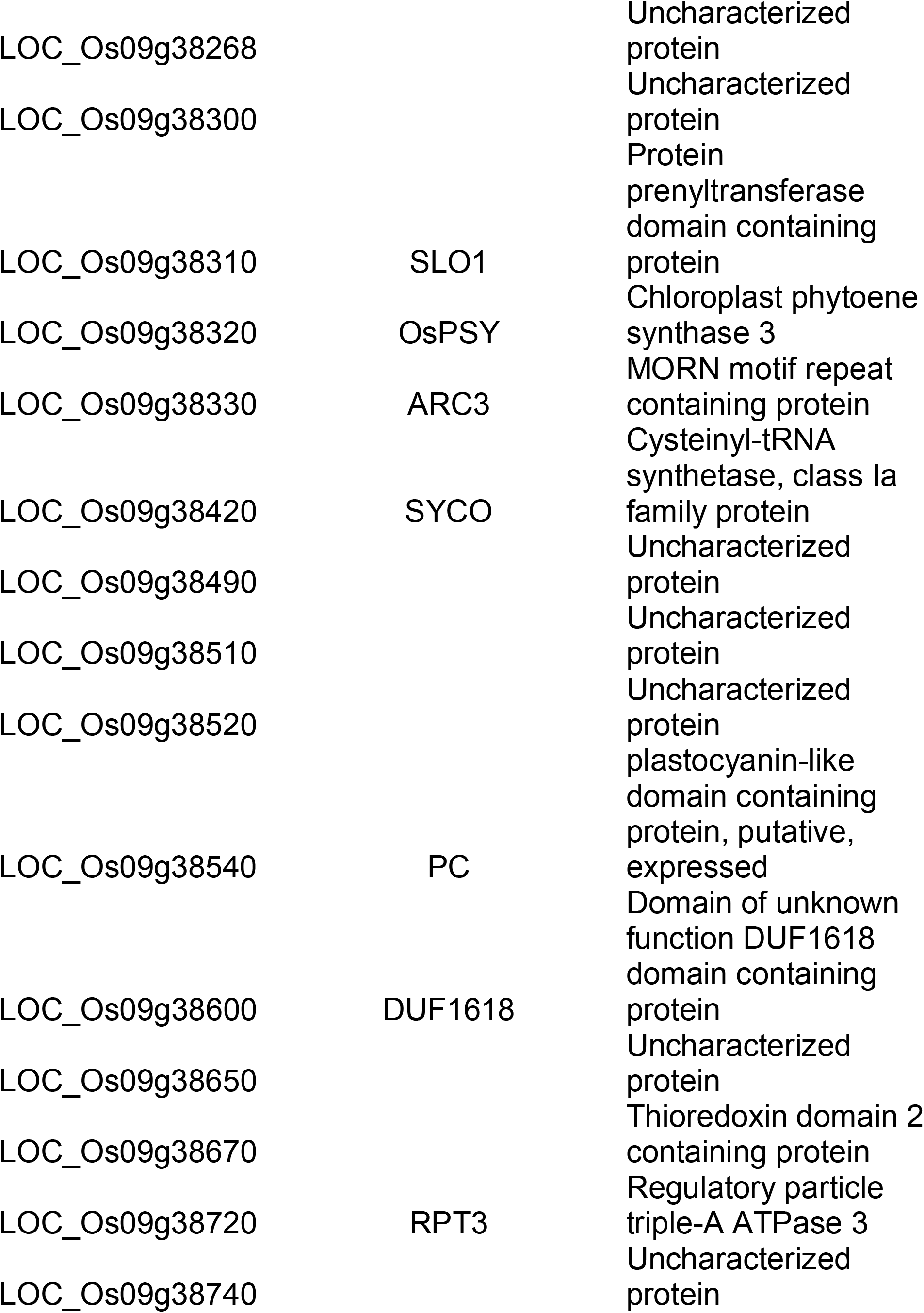
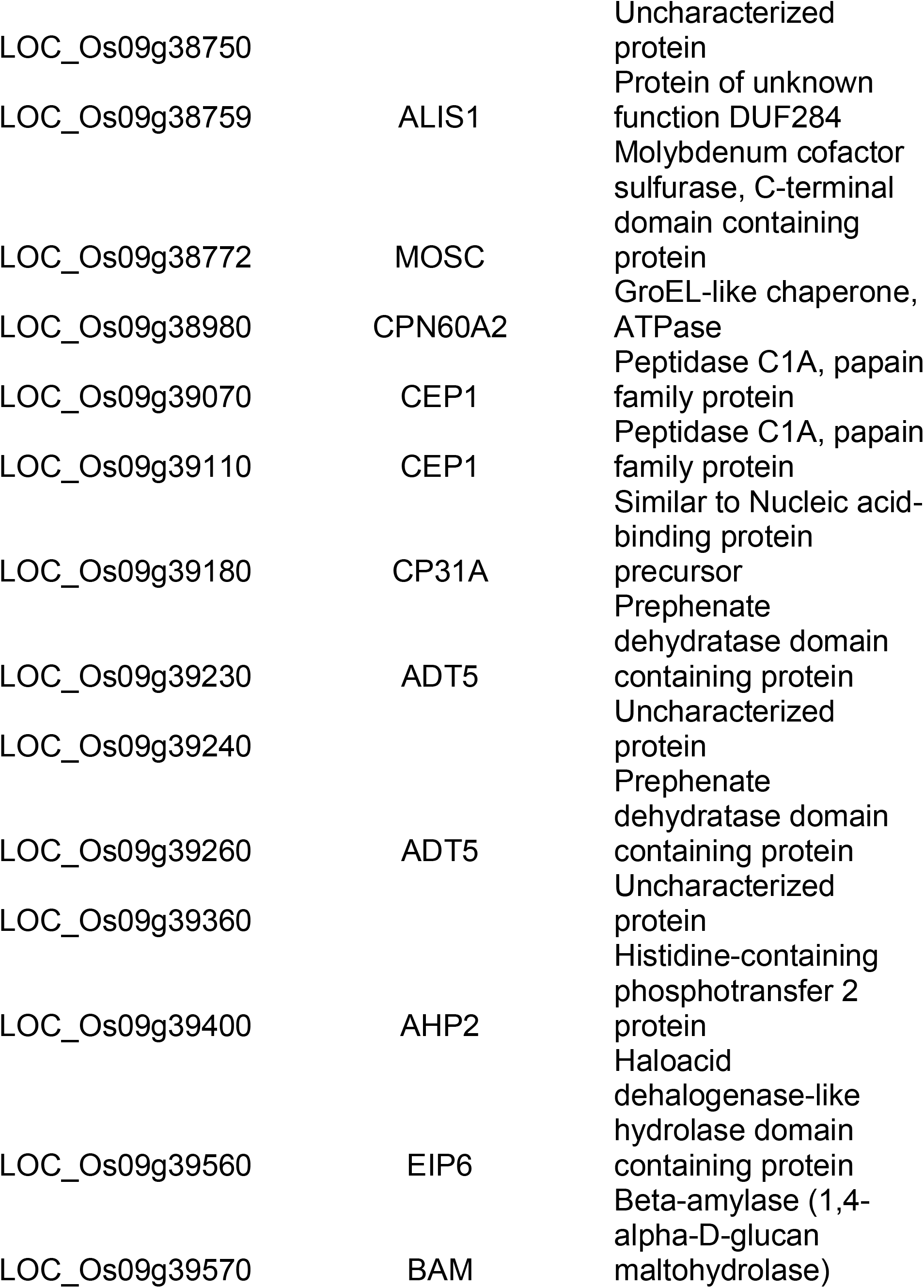
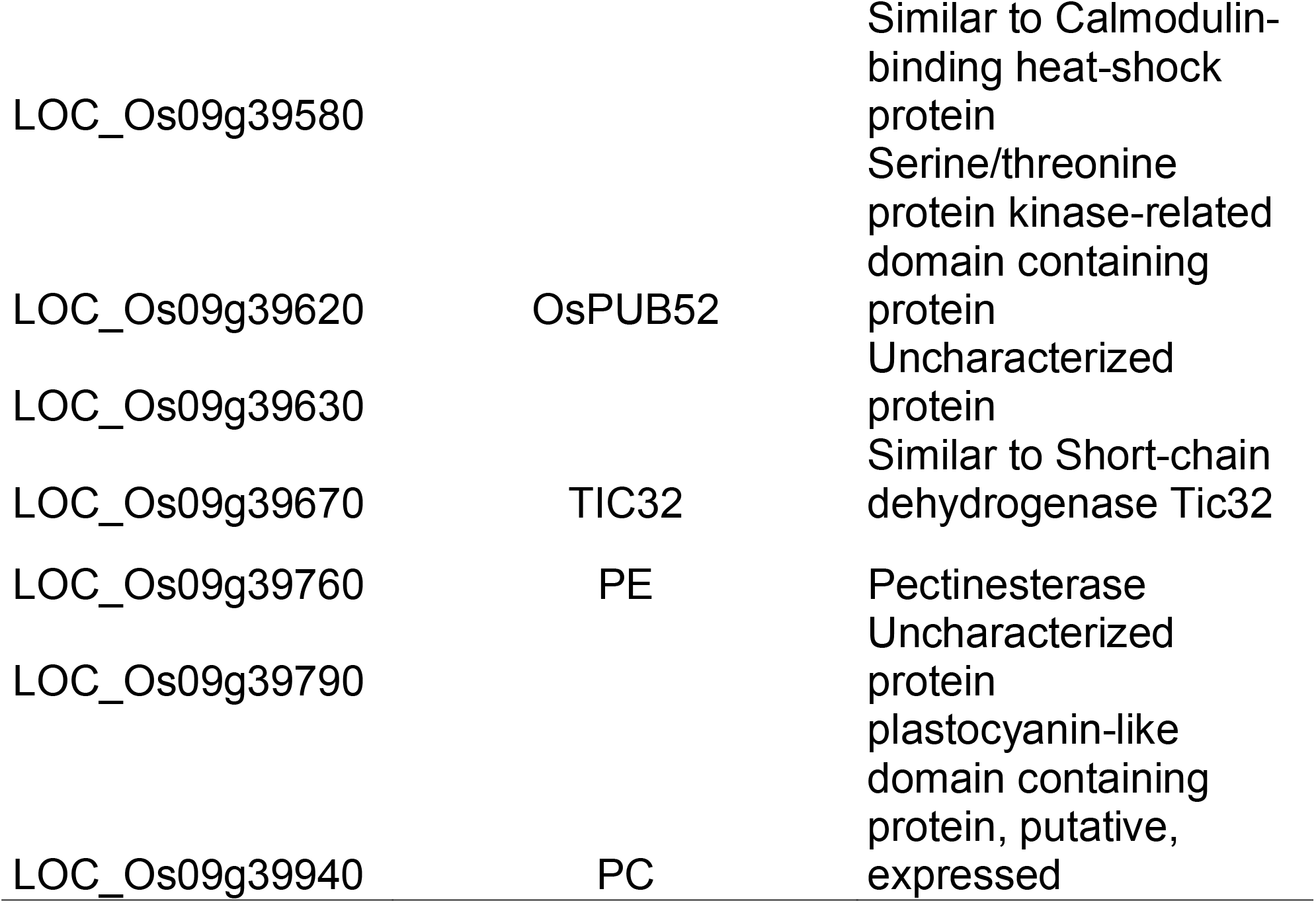
Description of genes from the GWAS peak on chromosome 9 of proteins with chloroplastic subcellular localization

### 3.3. Relationship between the haplotypes from the CGPI_ABS_, panicle weight and the rice population structure

The phenotypication in PI_ABS_ used in Experiment 2 was performed in RPD1 plants at tillering stage that posteriorly was used to determine the yield component parameters (YCP) panicle weight (PW) at the end of the grain filling stage. On the other hand, this has allowed the characterization of the haplotype distribution derived from the CGPI_ABS_ within the rice population structure. On the other hand, it has also allowed the characterization of the PW data distribution within the *O. sativa* subspecies and subpopulations and then to compare these with the haplotype distribution.

A PCA was performed using two sets of PI_ABS_ data: the PI_ABS_ average high (HP) and low (LP) values of each haplotype according to Fig. 3C and the particular PI_ABS_ of each genotype. This analysis indicated that the variability associated with all parameters agglomerated the data from HP and LP haplotypes in practically two isolated groups. This phenomenon is because the LP haplotypes only presented a few HP haplotype data, but the HP haplotypes did not group LP haplotype data (Fig. 4A). Another PCA was also performed using the previous two sets of PI_ABS_ data but identifying genotypes of the five *O. sativa* subpopulations. This PCA indicated that the group of HP haplotypes data were mainly composed of *Japonica* ssp. genotypes from the temperate-japonica, tropical-japonica and aromatic subpopulations (Fig. 4B).

**FIGURE 4.**
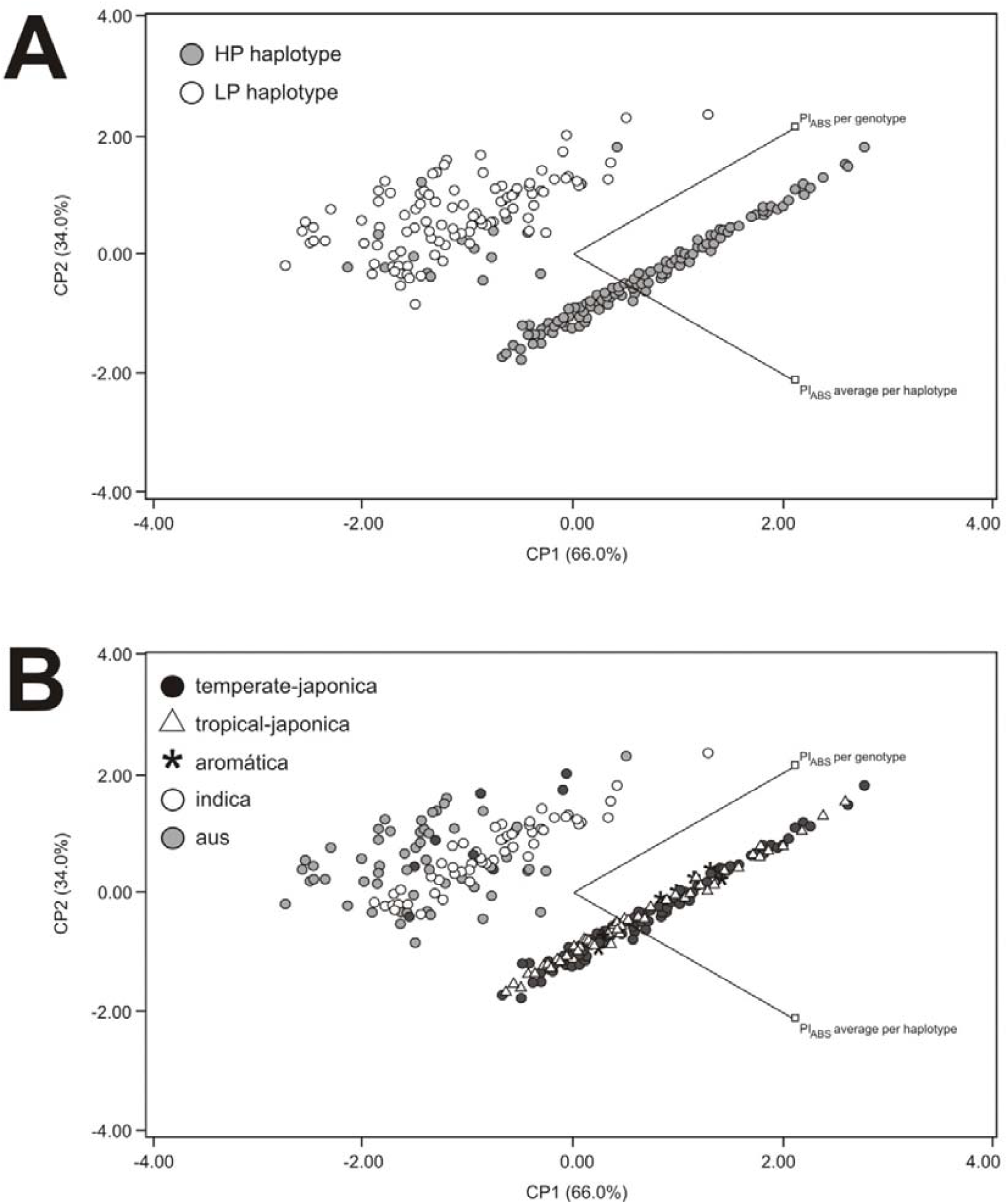
PCAs of PI_ABS_ average value per haplotype and PI_ABS_ per genotype data. HP and LP haplotypes associated with each CGPI_ABS_. **A**. the dots represent CP1 and CP2 values associated with HP and LP haplotypes. **B**. the dots represent CP1 and CP2 values associated with genotypes from each *O. sativa* subpopulation.

On the other hand, the group of LP haplotypes data was mainly composed of *Indica* ssp. genotypes from indica and aus subpopulations. Data could suggest that HP and LP haplotypes were related to *Japonica* spp. and *Indica* spp. subpopulations, respectively. However, these results also suggested a relationship between HP and LP haplotypes with PW because *Japonica* ssp. subpopulations presented higher PW values than *Indica* ssp. subpopulations (Fig. 5).

**FIGURE 5.**
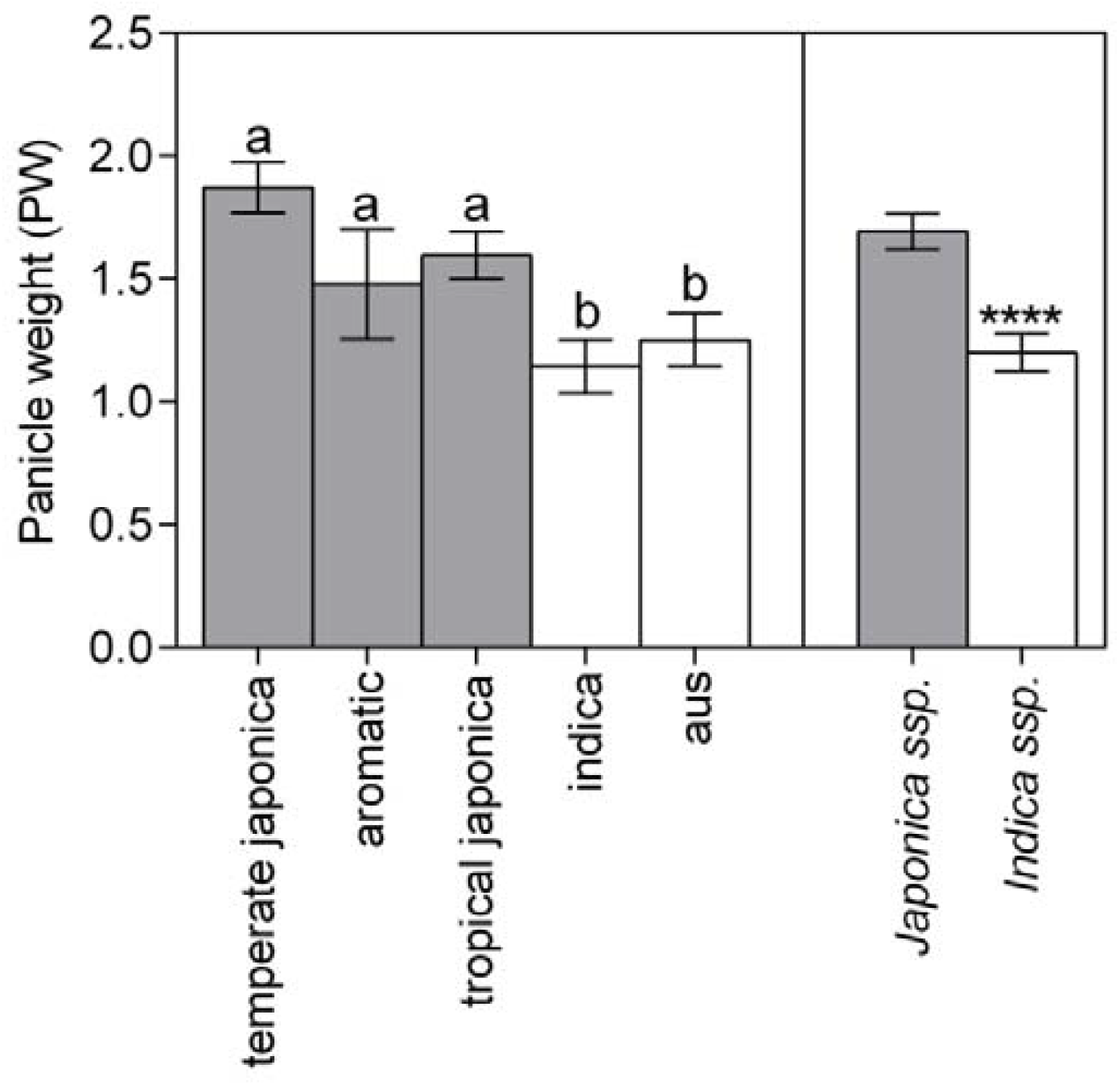
Panicle weight distribution within the *O. sativa* structure population. PW was determined in plants of the rice diversity panel RDP1 at the end of the filling grain. Gray (*Japonica* spp. genotypes) and white (*Indica* spp. genotypes) represent means ± S.E. Different letters represent significant differences between subpopulations (ANOVA and post hoc analysis DGC tests, *p* < 0.05, n = 3 per genotype). Asterisks represent significant differences between subspecies (Student’s t-test, two samples; \*\*\*\**p* < 0.0001; n = 3 per genotype).

## 4. Discussion

In the present work, we followed an approach based on different reports suggesting that plant screening by chlorophyll fluorescence analysis in flag leaf could be beneficial for breeding programs to improve grain productivity. The authors of these reports indicated that the OJIP parameter PI_ABS_ could be a robust indirect indicator of yield in rice (Zhang et al., 2015) but also in other crops such as wheat (Vuletić et al., 2019) and oat (Tobiasz-Salach et al., 2019). Our work supported these results suggesting that OJIP parameters determined in flag leaf, particularly PI_ABS_, are associated with grain yield parameters in rice. We have come to this conclusion after examining the statistic PCA, linear correlation analysis and linear regression analysis results that indicated a high positive correlation between all OJIP parameters, particularly with Y. However, PI_ABS_ explained a significant part of Y variation compared with the others OJIP parameters indicating that it could be the better indirect indicator of yield in rice. This data also suggested that the relationship between the photosynthetic activity and the grain yield would be explained by more than one particular structure or functionality of PSII, particularly those represented by PI_ABS_.

The PI_ABS_ has been analyzed as a phenotypic trait through GWAS techniques in different plant species studies (Liu et al., 2017; Chen et al., 2020; Zou et al., 2022) but it is only recently that some authors have published works in rice (Vilas et al., 2020; Khan et al., 2021). These last works indicated significant SNPs associated with PI_ABS_ and other OJIP parameters and that many SNPs, including those linked to PI_ABS_, were associated with genes related to the chloroplast. Notably, Vilas et al. (2020) indicated that some of these genes were CGPI_ABS_. Khan et al. (2021) also analyzed the PI_ABS_ distribution among *O. sativa* subpopulations indicating that the *Japonica* spp. subpopulations had higher PI_ABS_ values than *Indica* spp. subpopulations. However, needed to analyze the distribution of the HP and LP haplotypes and yield parameters among these subpopulations. Our analyses supported this result in terms of PI_ABS_ but, at the same time, suggested that this PI_ABS_ distribution was linked to the HP and LP genotypes distributions because these were associated mainly with the *Japonica* spp. and *Indica* spp. subpopulations, respectively. Even more, because our data indicated that the *Japonica* spp. subpopulations presented higher PW values than the *Indica* spp. subpopulations, this led us to think that some CGPIABS would explain the relationship between PI_ABS_ with the yield parameters. However, even more interesting, all this suggested that PI_ABS_ could be an early predictor of grain yield at the tillering stage in rice breeding processes.

74% of our characterized CGPI_ABS_ were annotated as a characterized protein. These protein characterizations were analyzed using hypothetical processes that can improve grain yield. Several CGPI_ABS_ were characterized as small auxin-up RNA (SAUR) proteins. The SAUR is a family of early auxin-responsive genes where almost 70% of its members were subcellular and located in chloroplast, including the SAURs in rice (OsSAURs) described by us (Zhang et al., 2021). Many SAUR proteins were associated with developmental processes in rice, particularly in reproductive development (Fujita et al., 2010; Courdet et al., 2011). The gene LOC_Os09g37690 encodes a flavin-containing monooxygenase (FMO) similar to FMO1 from Arabidopsis. The genes LOC_Os09g39230 and LOC_Os09g39260 encode arogenate dehydratase 5 (ADT5) from Arabidopsis. These last three genes were also related to the OsSAURs described above because they play an essential role in auxin biosynthesis (Zhao et al., 2001; Aoi et al., 2020). Some authors reported that the content of endogenous auxin in rice plants during the early maturing period is high (Dong et al., 2012), while other authors reported a PW increase in rice plants sprayed with auxin (Ahmadi and Nejad, 2014). Therefore, all this information led us to think that the set of CGPI_ABS_ related to auxin biosynthesis could influence the auxin level during panicle development. Consequently, they could influence the panicle weight, as was determined in Experiment 2.

Another considerable number of CGPI_ABS_ were related to the functional and structural quality of the chloroplast. Among these genes is found LOC_Os09g39570, which encodes a β-amylase involved in starch hydrolysis and some authors related it with panicle development in rice (Wang et al., 2019). Another three cases are the genes LOC_Os09g38320, LOC_Os09g39670 and LOC_Os09g38330, which encode a phytoene synthase, a short-chain dehydrogenase Tic32 and a protein ACCUMULATION AND REPLICATION OF CHLOROPLASTS 3 (ARC3), respectively. In the first case, the phytoene synthase is involved in the terpene synthesis necessary for chloroplast differentiation (Chaudhary et al., 2019). A gene encoding phytoene synthase was proposed in breeding strategies for improving rice yield using agrobacterium-mediated transformation (Khan et al., 2015). Tic32 and ARC3 are necessary at the chloroplast level because the first is an essential Component in Chloroplast Biogenesis (Hörmann et al., 2004), and the second was reported for its role in chloroplast division and expansion in *Arabidopsis* (Pyke and Leech, 1994). Also, genes LOC_Os09g38540, LOC_Os09g39940 and LOC_Os09g38980 encode proteins related to the biosynthesis of the component from the photosynthesis process. The first two genes encode plastocyanin, an electron transfer agent between cytochrome f of the cytochrome b6f complex from PSII and P700^+^ from PSI. The third gene encodes a chloroplastic-like chaperonin, which is an essential part of the system for folding the large subunit of ribulose 1,5-bisphosphate (Zhao and Liu, 2017). Several reports indicate that the flag leaf’s photosynthesis constitutes 60 to 100% of the carbonated structures allocated in rice grains (Wada et al., 1993; Takai et al., 2005). All these CGPI_ABS_ could influence the photosynthetic apparatus performance, which could consequently act indirectly on yield parameters.

A common feature of almost all CGPI_ABS_ described in this work was that they were reported as candidate genes in different studies related to rice yield breeding in different situations. For example, the gene LOC_Os09g37610 was recently identified in rice breeding for grain-balanced elemental concentrations (Dwiningsih and Alkahtani, 2022). The genes LOC_Os09g39070 and LOC_Os09g39240 were linked with iron uptake in roots (Weirich et al., 2019). Many others as LOC_Os09g38420, LOC_Os09g38510, LOC_Os09g39070, LOC_Os09g39180, LOC_Os09g39620, LOC_Os09g39670, and LOC_Os09g39760, were reported in breeding studies to improve rice for different abiotic and biotic tolerance (Wang et al., 2014; Du et al., 2015; Raorane et al., 2015; Lakra et al., 2018; Gongora, 2015; Cal et al., 2019). Our results suggested that all these genes could be candidates for use as molecular markers in a molecular assistance process for rice breeding for grain productivity in standard field conditions. In this regard, only the candidate gene LOC_Os09g39240 has been reported by a few authors for its relationship with rice breeding for grain productivity under this condition (Ramos et al., 2019).

## 5. Conclusions

The challenge of improving rice plants for grain productivity involves developing efficient tools that accelerate the plant selection process through the rapid determination of robust indicators that allow the analysis of a large number of plants. If such determinations involve a low cost per data point, the plant selection process would have the potential to make scalable any breeding program. Here we have found associations between grain yield and OJIP parameters determined in the flag leaf, among which the PI_ABS_ stands out for its strong association with the yield. Early phenotyping by determining PI_ABS_ in plants from a rice diversity panel led to the characterization of a QTL with a set of various candidate chloroplastic genes with contrasting haplotypes for PI_ABS_. Combined analysis of these haplotypes in terms of population structure in rice indicated that these were distributed within the subpopulations of *O. sativa* similarly to the population structures described to PW. This data suggested that the PI_ABS_ determination could be an early predictor of yield in rice breeding processes. At the same time, the haplotypes of the candidate genes associated with PI_ABS_ could also be used to perform molecular assistance in this process. However, further studies are necessary to determine their effectiveness.

## Abbreviations

GWAS: genome-wide association analysis
LD: linkage disequilibrium
MAF: minor allele frequency
PW: panicle weight
PI_ABS_: photosynthetic performance index
PCA: principal component analyses
QTL: quantitative trait locus
RC: active PSII reaction centre
RDP1: Rice Diversity Panel 1
SNPs: single nucleotide polymorphism
WFG: weight of filled grains
Y: Kg of grains per ha^-1^

## 6. Acknowledgements

This work was financially supported by projects PICT 2019-02779, funded by Agencia Nacional de Promoción Científica y Tecnológica, Argentina and PIP0363, funded by Comision Nacional de Investigaciones Científicas y Técnicas.

