## Supplemental Table 1 for "Field and genetic evidence support the photosynthetic performance index (PI_ABS_) as an indicator of rice grain yield"

| Variables | e1 | e2 |
| --- | --- | --- |
| Kg . m^-2^ | 0.42 | 0.82 |
| grains . m^-2^ | 0.29 | 0.12 |
| PI_ABS_ | 0.45 | -0.18 |
| ΨE_O_ | 0.44 | -0.00 |
| F_V_/F_M_ | 0.43 | -0.24 |
| γRC | 0.40 | -0.40 |

**Table S1. Eigenvectors associated to each original variable.** Eigenvectors associated to each original variable weighted to form PC1 (e1) and PC2 (e2).
